# Development of non-transgenic glyphosate tolerant wheat by TILLING

**DOI:** 10.1101/2020.07.23.218883

**Authors:** Charles P. Moehs, William J. Austill, Daniel Facciotti, Aaron Holm, Dayna Loeffler, Zhongjin Lu, Jessica C. Mullenberg, Ann J. Slade, Michael N. Steine, Jos van Boxtel, Cate McGuire

## Abstract

Glyphosate (N-phosphonomethyl-glycine) is the world’s most widely used broad spectrum, post-emergence herbicide. It inhibits the chloroplast-targeted enzyme 5-enolpyruvylshikimate-3-phosphate synthase (EPSPS; EC 2.5.1.19), a component of the plant and microorganism-specific shikimate pathway and a key catalyst in the production of aromatic amino acids. Variants of EPSPS that are not inhibited by glyphosate due to particular amino acid alterations in the active site of the enzyme are known. Some of these variants have been identified in weed species that have developed resistance to glyphosate because of the strong selective pressure of continuous, heavy glyphosate use. We have used TILLING (Targeting Induced Local Lesions in Genomes), a non-transgenic, target-selected, reverse genetics mutation breeding technique, and conventional genetic crosses, to identify and combine, through two rounds of mutagenesis, wheat lines having both T_102_I and P_106_S (so-called TIPS enzyme) mutations in both the A and the D sub-genome homoeologous copies of the wheat EPSPS gene. The combined effects of the T_102_I and P_106_S mutations are known from previous work in multiple species to minimize the binding of the herbicide while maintaining the affinity of the catalytic site for its native substrates. These novel wheat lines exhibit substantial tolerance to commercially relevant levels of glyphosate.

## Introduction

Glyphosate (N-phosphonomethy-glycine) is the world’s most widely used herbicide (Duke and Powles 2008) (Duke et al. 2018). It forms the basis of the commercial Roundup^®^ ready system in which multiple crops such as maize (*Zea mays* L.), soybeans [*Glycine max* (L.) Merr.], cotton (*Gossypium hirsutum* L.), canola (*Brassica napa* L.), alfalfa (*Medicago sativa* L.) and sugar beets (*Beta vulgaris* L.) have been rendered resistant to the effects of glyphosate-containing herbicides by genetic transformation with herbicide insensitive forms of the glyphosate target enzyme 5-enolpyruvylshikimate-3-phosphate synthase (EPSPS) (Green 2018). This enzyme is the penultimate enzyme of the shikimate pathway, which in plants, bacteria and fungi is responsible for the synthesis of aromatic amino acids and downstream metabolites (Tzin and Galili 2010). In plants, glyphosate is rapidly translocated to the meristems and kills these growing points by starving them of these essential amino acids (Duke et al. 2003).

Due to the strong selection pressure of continuous, heavy use of this herbicide in glyphosate-resistant crops, many weeds have developed resistance to glyphosate and multiple resistance mechanisms are known (Heap and Duke 2018). These include target site mutations in the EPSPS enzyme itself, as well as other mechanisms including gene amplification of EPSPS (Gaines et al. 2010) (Gaines et al. 2019) and mutations that increase vacuolar sequestration of glyphosate (Ge et al. 2014). In some cases, weed species exhibit combinations of multiple resistance mechanisms (Gherekhloo et al. 2017) (Sammons and Gaines 2014).

Among the target site mutations identified to date, missense mutations at amino acid 106-the numbering is based on the mature form of the *Arabidopsis* EPSPS-are most frequently found. The normally occurring proline at this position is altered to serine, threonine, valine and leucine in various glyphosate tolerant weed species (reviewed in: (Gaines et al. 2020)). These single missense mutations impart a 2-6 fold increase in glyphosate tolerance. Greater glyphosate tolerance has evolved in goosegrass (*Eleusine indica*) with the combination of the P_106_S missense alteration with a T_102_I mutation (known as the TIPS enzyme) (Yu et al. 2015), which leads to an EPSPS enzyme that is essentially insensitive to the effects of glyphosate (Funke et al. 2009). However, the TIPS enzyme in goosegrass is associated with a fitness cost due to its lowered catalytic efficiency (Han et al. 2017) (Vila-Aiub et al. 2019). Other double mutant and even a triple mutant EPSPS have evolved in response to herbicide pressure (Perotti et al. 2019) (Takano et al. 2020).

Glyphosate resistant wheat was generated transgenically (Zhou et al. 2003), but, due to opposition by growers and consumers, was never commercialized or widely grown (Dill 2005). Nevertheless, a glyphosate-resistant wheat would offer benefits to growers in some geographies for post-emergent control of otherwise difficult to control weeds, including wild oats (*Avena fatua* L.), feral rye (*Secale cereal* L.), jointed goat grass (*Aegilops cylindrica*), downy brome (*Bromus tectorum* L.) and blackgrass (*Alopecurus myosuroides*). These grass weeds can be difficult to control with currently available options and impose considerable economic costs. For example, in the United Kingdom, it was recently estimated that blackgrass control, and the yield losses in winter wheat that it causes, may result in an annual economic burden as high as £1 billion (Varah et al. 2019). These issues have prompted efforts to develop wheat resistant to glyphosate (Aramrak et al. 2018) and other herbicides (Newhouse et al. 1992) (Ostlie et al. 2015) (Pozniak et al. 2004) via non-transgenic (mutagenesis) methods that might be more acceptable to growers and the public. These efforts have had some success with the development of wheat lines with some glyphosate tolerance and wheat with tolerance to other herbicides through forward screening of mutagenized wheat seeds. However, the genetic basis of wheat glyphosate tolerance in the described glyphosate tolerant lines is incompletely understood (Aramrak et al. 2018) and the level of glyphosate tolerance is insufficient to be commercially useful.

Here, we describe our use of the non-transgenic method of TILLING (Colbert et al. 2001) (McCallum et al. 2000) (Slade et al. 2005) to create and identify non-transgenic target site mutations in the A, B and D sub-genome homoeologous copies of the wheat EPSPS gene. Through two rounds of mutagenesis, we were able to identify wheat lines containing both active site T_102_I and P_106_S mutations in the A and, separately, in the D genome copy of EPSPS (in wheat these amino acid positions are equivalent to T_168_I and P_172_S using the full-length wheat 7A EPSPS protein as reference (KP411547, (Aramrak et al. 2015), but for simplicity’s sake we will use the conventional numbering based on the mature *Arabidopsis* or maize EPSPS sequence (CAA44974.1, (Dong et al. 2019)). We created lines containing both A and D sub-genome homozygous copies of this double mutant combination through the use of conventional genetic crosses and these lines exhibited substantial tolerance to commercially relevant levels of glyphosate in the greenhouse and in the field.

## Results

### Identification of novel EPSPS TILLING alleles and glyphosate resistant phenotype

At the commencement of this project, the wheat genome sequence was not available and only cDNA and EST sequences putatively encoding wheat EPSPS homoeologs were available in public databases. We queried available wheat nucleotide resources, including genomic and EST databases at NCBI, with the known rice EPSPS genomic and cDNA sequences (Junwang et al. 2002) to identify homologous sequences in wheat. The ESTs were downloaded from NCBI and assembled using DNASTAR Lasergene software. On the assumption that the intron-exon structure of the rice EPSPS gene and the wheat gene are similar, we designed primers near the likely N-terminal region of wheat EPSPS exon 2 and sequences expected to be near the C-termini of wheat EPSPS exons 4 and 5 (**Figure 1A**). Since the known conserved active site region of the wheat EPSPS gene, like the rice EPSPS gene, was likely to be present near the C-terminal end of the 2^nd^ exon, the primer pairs we designed were intended to encompass this active site region. We used these primers to amplify PCR bands from genomic DNA of hexaploid bread wheat (cultivar Express) and these bands were cloned and sequenced. The sequenced bands fell into three sequence classes that putatively arose from an EPSPS gene on each of the three homoeologous genomes of hexaploid wheat. The sequences exhibited SNPs near the active site of the enzyme that clearly distinguished the different homoeologous copies of the gene (**Figure 1C**). Isolation of RNA from leaf tissue and sequencing of cDNA with EPSPS-specific primers identified mRNA fragments homologous in sequence to each of the three homoeologs, indicating that each of these homoeologs is expressed. We did not find evidence for expression of more than three copies of EPSPS. Based on the sequences of the different SNP-containing EPSPS gene fragments, we designed new primer pairs that specifically amplified each of the individual homoeologous sequences (**Table 1**, **Figure 1B**). These new homoeolog-specific primers exploited sequence differences, principally in the introns, that distinguished the homoeologs. Additionally, we confirmed that the different genomic sequences we had isolated arose from different homoeologs by amplifying these fragments in chromosome deletion lines of wheat. This determined that EPSPS sequence variant A is present on wheat chromosome 7A, variant B is encoded by chromosome 4A-due to a translocation (Chao et al. 1989), and variant D is present on homoeologous chromosome 7D (**Figure 1C**). The chromosome assignments of these sequences has been determined by others as well (Aramrak et al. 2015). Next, the identified primer pairs that specifically amplified single homoeologs were used to screen an existing TILLING resource of wheat cultivar Express (Slade et al. 2005) to identify novel SNPs in each of the wheat EPSPS homoeologs.

**Table 1.**
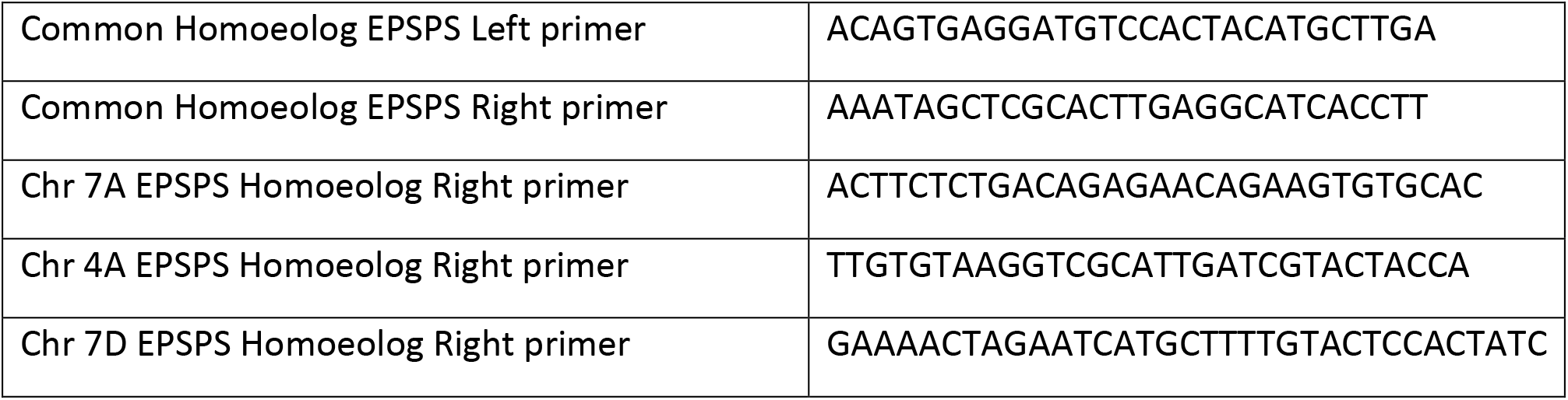
Primer pairs used for gene amplification, TILLING and genotyping.

**Figure 1.**
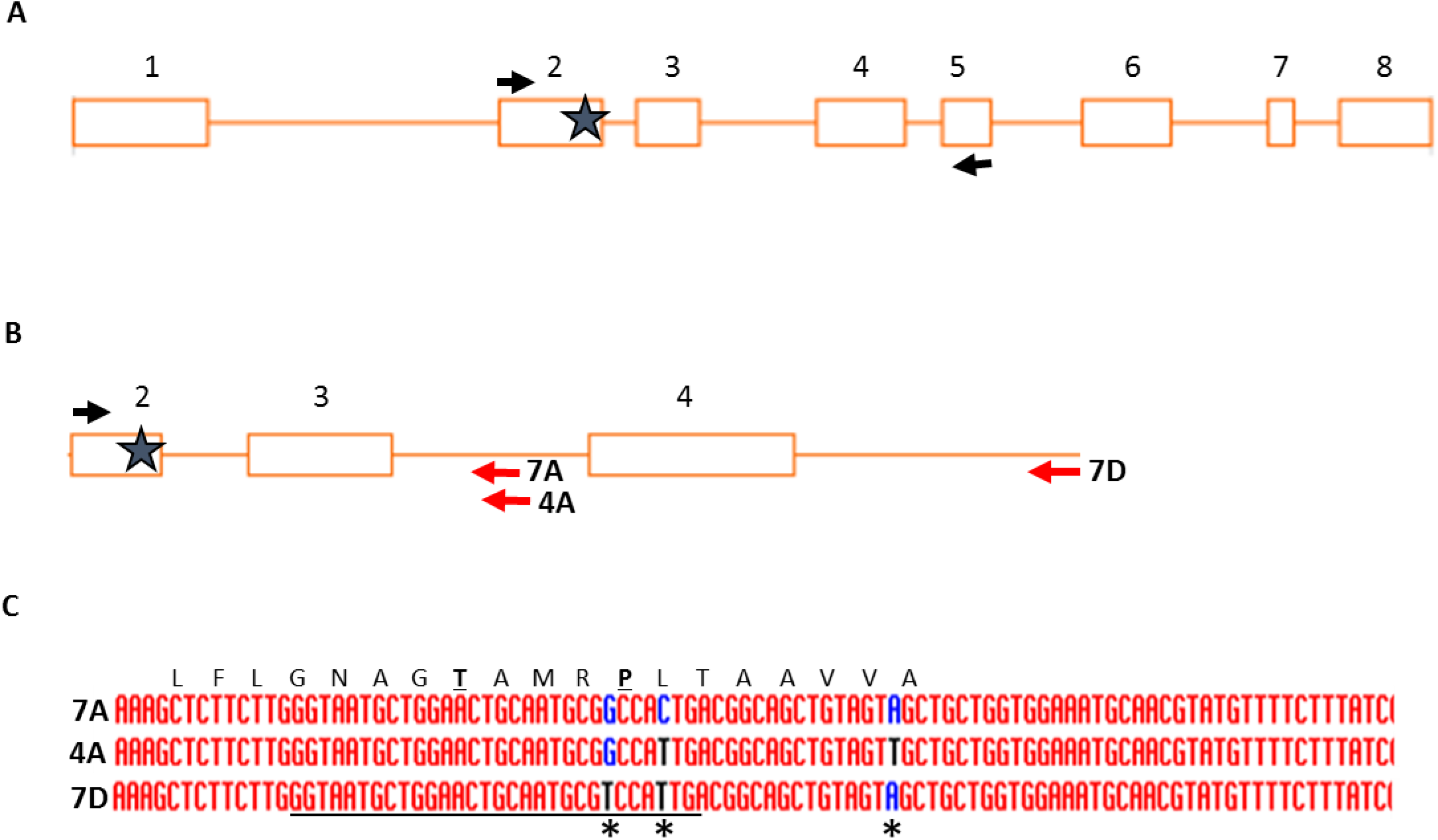
Overview of wheat EPSPS homoeolog cloning and TILLING strategy and homoeolog-specific SNPS. **A.** The genomic intron-exon structure of the rice EPSPS is shown. Boxes indicate the eight exons and lines represent introns. The black arrows indicate the locations of PCR primers used to amplify fragments of wheat EPSPS homoeologs from wheat genomic DNA spanning the active site of the enzyme-indicated with a star. **B.** TILLING of EPSPS homoeologs was accomplished with a common left primer (black arrow) and homoeolog-specific right primers (red arrows). **C.** Alignment of the wheat EPSPS homoeologs showing homoeolog-distinguishing SNPs-indicated with asterisks-surrounding the active site of the enzyme. Key active site residues threonine and proline are underlined. Alignment created using default parameters of the software: MultAlin (http://multalin.toulouse.inra.fr/multalin/) (Corpet 1988).

**Figure 2** highlights the priority TILLING alleles identified in the active site of the enzyme in the first round of screening (underlined: P_106_S in the 7A genome and T_102_I in the 7D genome). These alleles represent just two of the more than 50 alleles identified across the fragments screened. Because of previously published information about missense mutations that render EPSPS resistant to the effects of glyphosate, we focused our search for alleles on amino acids threonine 102 and proline 106. In the *E. coli* homolog of EPSPS, these amino acids are at positions 97 and 101 (Funke et al. 2009). The combination of two mutations at these sites, T_102_I and P_106_S, leads to an EPSPS variant referred to as the TIPS enzyme that is insensitive to the effects of glyphosate (Funke et al. 2009) (Sidhu et al. 2000) (Yu et al. 2015). Each of these individual mutations also affects the enzymatic properties of EPSPS. Novel TILLING alleles that were not near the active site of the EPSPS enzyme were not characterized further. The T_102_I allele in the D genome was the first priority allele identified, followed by the P_106_S allele in the A genome. Subsequently, we also identified a T_102_I allele in the EPSPS homoeolog on chromosome 4A. In each case, the identified mutant was heterozygous, which allowed us to identify segregating sibling homozygous mutant and wild type EPSPS plants in the M3 generation using KASP (Kompetitive allele specific PCR; https://www.biosearchtech.com/support/education/kasp-genotyping-reagents) primers developed to distinguish them.

**Figure 2.**
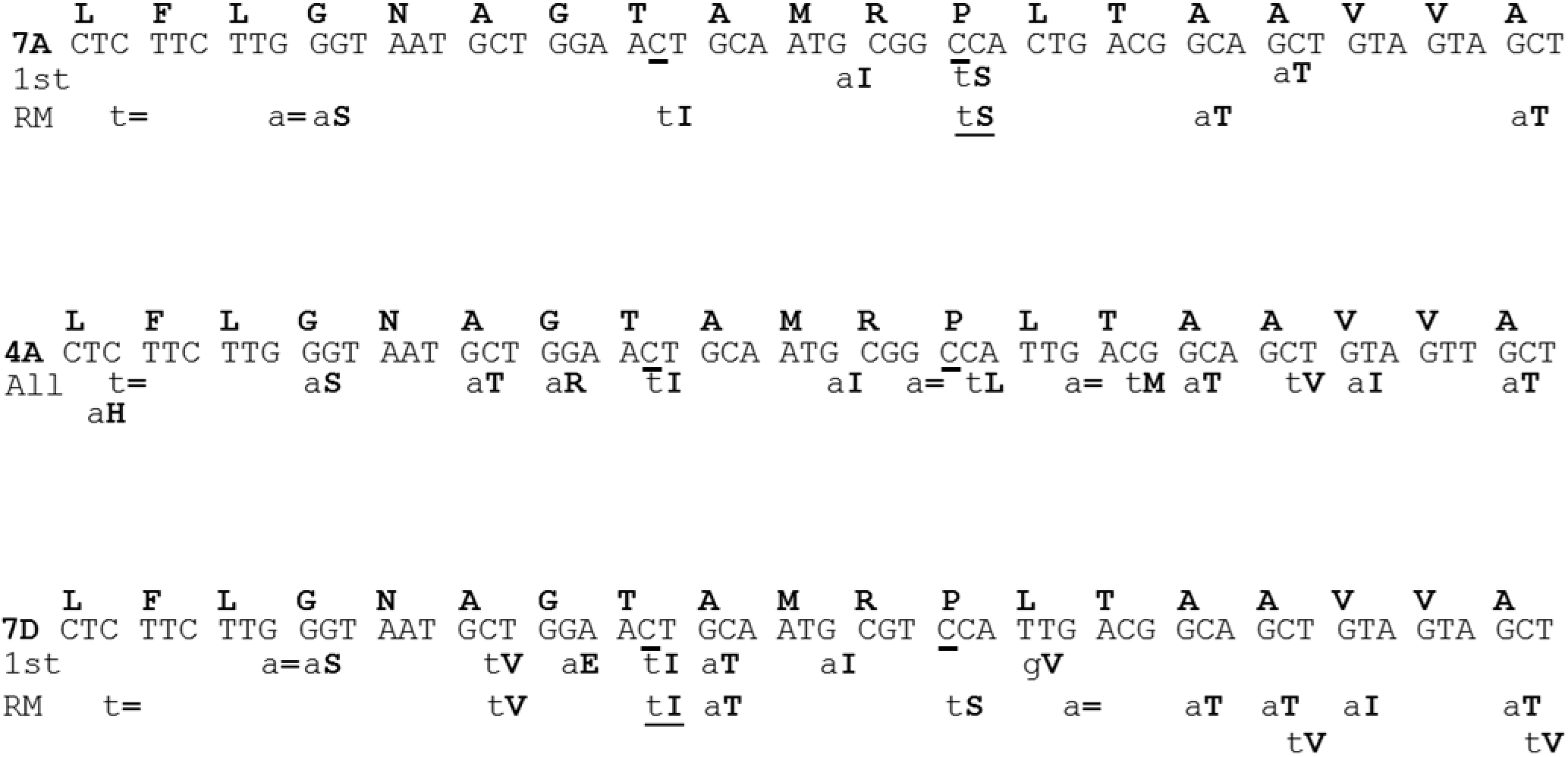
Depiction of TILLING alleles identified in the active site regions of wheat EPSPS homoeologs. Conserved regions around the active site of wheat 7A, 4A and 7D EPSPS homoeologs are depicted with identified novel induced SNPs shown below the homoeolog-specific DNA sequence. The 1^st^ row refers to the TILLING screen of the initial mutagenized population. The second row labeled “RM” refers to alleles identified upon remutagenesis. The nucleotides mutated to create the “TIPS” alleles are underlined. Lower case letters indicate the altered nucleotide while the capitalized bolded letters refer to the the amino acid alteration due to the SNP. Since the line that was remutagenized contained the 7A P_106_S and the 7D T_102_I alleles, these alleles are repeated in the “RM” row to indicate their presence in the remutagenized 2^nd^ TILLING population. All of the TILLING mutants identified in the 4A EPSPS homoeolog in both TILLING screens are shown on one row. The equal sign indicates a silent SNP that doesn’t alter the amino acid.

Homozygous lines containing each of the single A, B and D mutations as well as combinations containing two or three of the single mutations combined through conventional crosses were created (schematically illustrated in **Figure 3**) and initial glyphosate tolerance tests were conducted. Seeds homozygous for individual and combined priority alleles were germinated on glyphosate-containing medium and their ability to germinate and grow on this medium was determined (**Figure 4**). Subsequently, these lines were also grown in the field and sprayed with several levels of glyphosate-containing herbicide. The results indicated that these lines containing priority alleles could survive and continue to grow on medium, and set seed in the field, on levels of glyphosate and glyphosate-containing herbicide that were lethal to un-mutagenized control parental cultivar Express as well as to wild type sibling lines. Nevertheless, it was apparent that the level of glyphosate tolerance was insufficient to be commercially useful. This led us to seek additional mutants that would increase glyphosate tolerance further, by re-mutagenizing the homozygous line containing the 7A and 7D homoelogous mutant EPSPS alleles (**Figure 3B,** line **A**).

**Figure 3.**
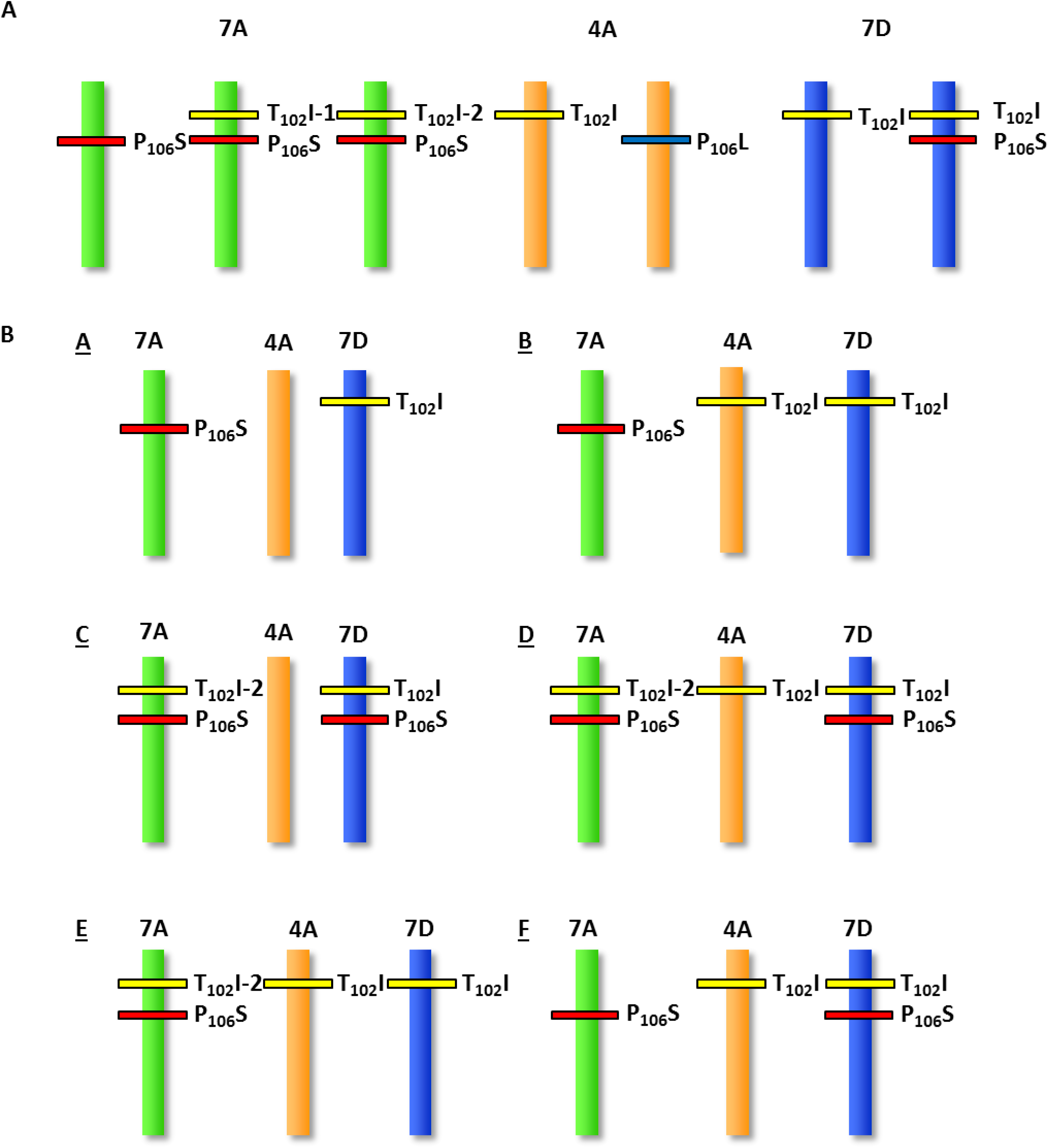
Schematic illustration of priority alleles identified and genetic constitution of lines created following conventional crosses. **A.** Priority alleles identified in the three targeted EPSPS homoeologs. The T_102_I allele on chromosome 7A was identified two independent times. **B.** Genetic constitution of wheat lines generated by conventional crosses. One chromosome is schematically illustrated but all lines are homozygous for the indicated alleles. (Not all possible combinations are shown.)

**Figure 4.**
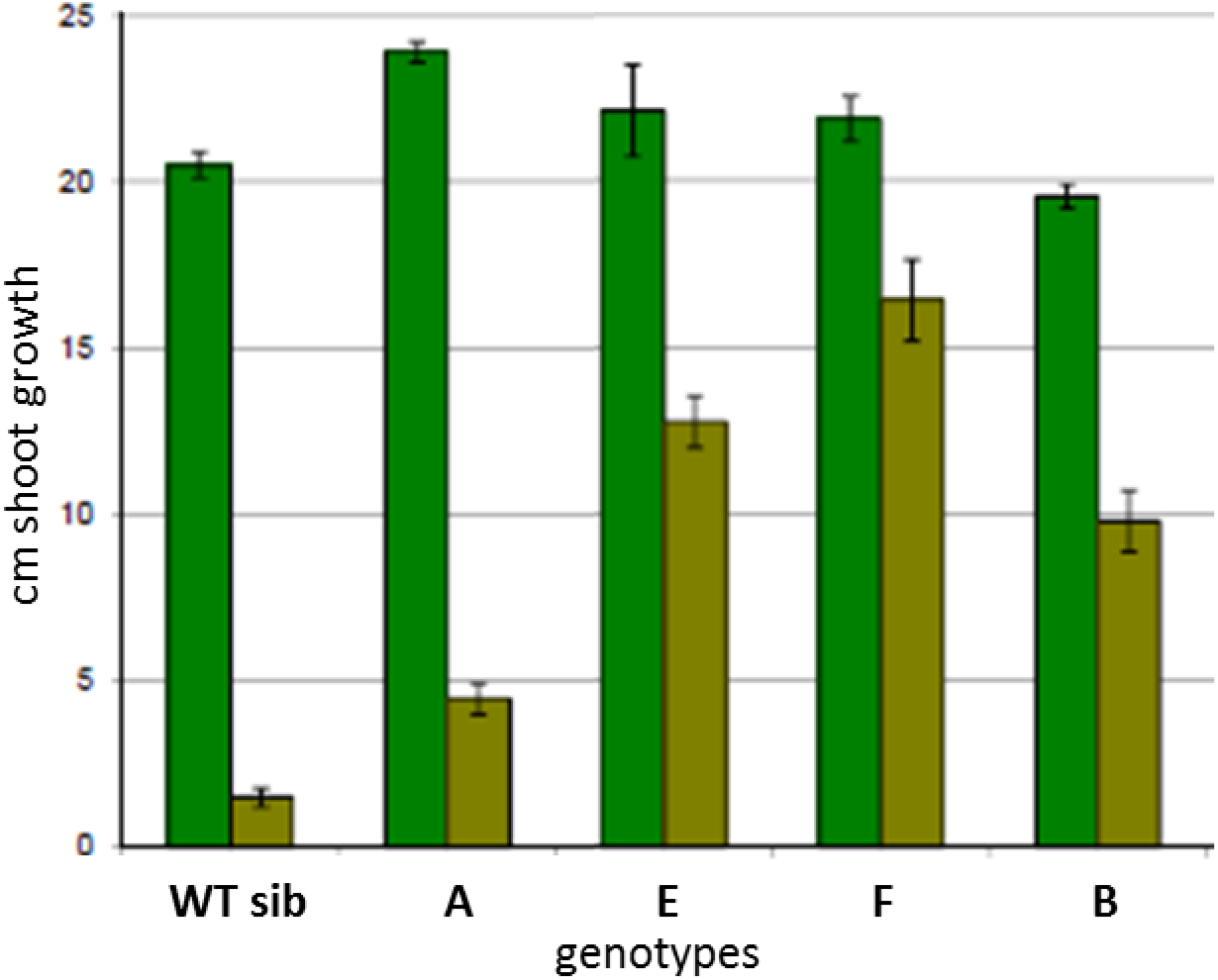
Shoot growth of different genotypes on Phytagel medium lacking or containing glyphosate. Average shoot height after twelve days is shown. Dark green bars represent medium without glyphosate. Light green bars indicate medium with 0.15mM glyphosate incorporated. Average +/− SE of six biological replicates is shown. Legend: **WT sib**: a wild type sibling line of a triple mutant line (genotype **B** in **Figure 3B**). **A** represents a double mutant line (genotype **A** in **Figure 3B**). **E** represents a line containing TIPS mutations on chromosome 7A and T_102_I mutations on both 4A and 7D chromosomes (genotype **E** in **Figure 3B**). **F** has the TIPS mutation on chromosome 7D and P_106_S on the 7A chromosome as well as the T_102_I mutation on chromosome 4A (genotype **F** in **Figure 3B**). **B** represents a genotype containing three single mutations (genotype **B** in **Figure 3B**).

### Re-mutagenesis, identification of additional EPSPS TILLING alleles and genetic crosses

The D genome T_102_I mutant EPSPS-containing line was crossed to the un-mutagenized parental variety, Express, and M2 mutant homozygotes were crossed to M3 homozygous progeny of the A genome P_106_S EPSPS mutant line. To generate a large amount of seed from A and D mutant EPSPS-containing homozygous progeny, M2 double homozygotes were identified, M3 seed was planted in the field and M4 seed was harvested. This M4 seed was used to generate a second (doubly mutagenized) TILLING population of approximately 10,000 individual M1 and an equivalent number of M2 members. Screening this population for additional alleles in the A and D genome copies of EPSPS led to the identification of new alleles: we identified an individual plant that in the background of the 7D genome T_102_I mutation also contained the nearby (11 nt away) P_106_S mutation. In addition, we identified two independent instances of plants containing the same TIPS mutant combination in the 7A genome copy of EPSPS. A plant containing a P_106_L mutation in the EPSPS homoeolog on chromosome 4A was also identified. An M4 plant homozygous for the 2^nd^ independently identified TIPS EPSPS chromosome 7A mutant combination was crossed to a plant homozygous for the three single mutants (7A_P_106_S; 4A_T_102_I; 7D_T_102_I) in order to also introduce the chromosome 4A genome mutant. The TIPS chromosome 7D mutant combination was likewise crossed in the same manner as the TIPS 7A mutant to the above triple mutant. Finally, in order to create plants containing the TIPS mutants on both chromosome 7A and 7D EPSPS genes, as well as plants that also contained a chromosome 4A T_102_I mutant, these two F1 plants, one heterozygous for the TIPS 7A mutants, and the other heterozygous for the TIPS 7D mutants were crossed to each other. F1 seeds from this cross were planted and genotyped and plants heterozygous for both 7A and 7D TIPS mutants were crossed to parental Express to reduce unwanted background mutations in these TILLING lines. This process of removing background mutations is ongoing and is currently at the BC3 generation. Crosses to remove background mutations were made between F1s rather than F2s to save a generation and speed this process. A schematic illustration depicting the principal priority alleles found and the genotypes of the lines that were created is shown in **Figure 3**.

### Assessment of glyphosate tolerance of the identified alleles and allele combinations

All of the lines containing single A, single D or single B genome mutant alleles, as well as their wild type sibling lines were first screened *in vitro* on glyphosate-containing phytagel medium before being grown in the field along with their EPSPS wild type siblings and un-mutagenized Express cultivar control. These experiments revealed that, individually and combined, the EPSPS mutant seeds-whether A genome P_106_S or D genome T_102_I-survived and grew when plated on low levels (0.15mM) of glyphosate-containing phytagel medium, including the growth of roots directly into the medium (not shown), while the wild type siblings did not survive (**Figure 4**).

The double and triple single mutation lines (A and B in **Figure 3B**) were also grown in a field trial. This trial revealed that although these mutations enabled the plants to survive doses of glyphosate herbicide (16 fl oz./acre) that were lethal to un-mutagenized parental control cultivar Express and to wild type (at the EPSPS locus) siblings of the mutants, the seed yield of the mutants was depressed in comparison to the mutants unsprayed by the herbicide. In addition, the mutants exhibited a yield depression, in the absence of herbicide treatment, compared to the parental cultivar, presumably due to the general effects of mutagenesis treatments. In our experience with a range of TILLING mutants in other genes, these mutagenesis treatment effects can be largely eliminated with several crosses to unmutagenized material.

The results of the *in vitro* and field experiments to judge the glyphosate tolerance of the first single mutant EPSPS combinations determined that these mutants did not confer sufficient glyphosate tolerance to the plants to be commercially useful. As the doubly mutagenized TILLING resource became available and the 7A and 7D genome TIPS mutants were discovered, we repeated the previous phenotyping experiments on these new mutants and their combinations. *In vitro* experiments (**Figure 4, E** and **F**) indicated that TIPS mutant-containing lines showed more robust survival and root extension into the medium than the previously tested single mutants. Spray chamber tests (**Figure 5, C** and **D**) and field growth (**Figure 6**) of the homozygous 7A and 7D double TIPS mutant lines revealed substantially enhanced herbicide tolerance compared to the single mutant lines. The double TIPS mutant was able to withstand the effects of a 28 fl oz./acre Roundup PowerMAX treatment, a treatment that led to complete mortality of the parental Express cultivar. Almost complete mortality of parental Express was observed at 11 fl. oz/acre, and at 16 fl. oz/acre no parental Express wheat survived. Although the TIPS lines subsequently set seed in the field, the seed yield was approximately 90% of the untreated TIPS line. Each of the three yield components: number of heads/area, number of seeds/head and 1000 seed weight were reduced approximately equivalently due to glyphosate treatment: between 5-14% each. However, these lines also exhibited reduced yield due to the mutagenesis treatment. Therefore, it is premature to assess the ultimate level of glyphosate tolerance achieved until removal of unwanted background mutations is further advanced.

**Figure 5.**
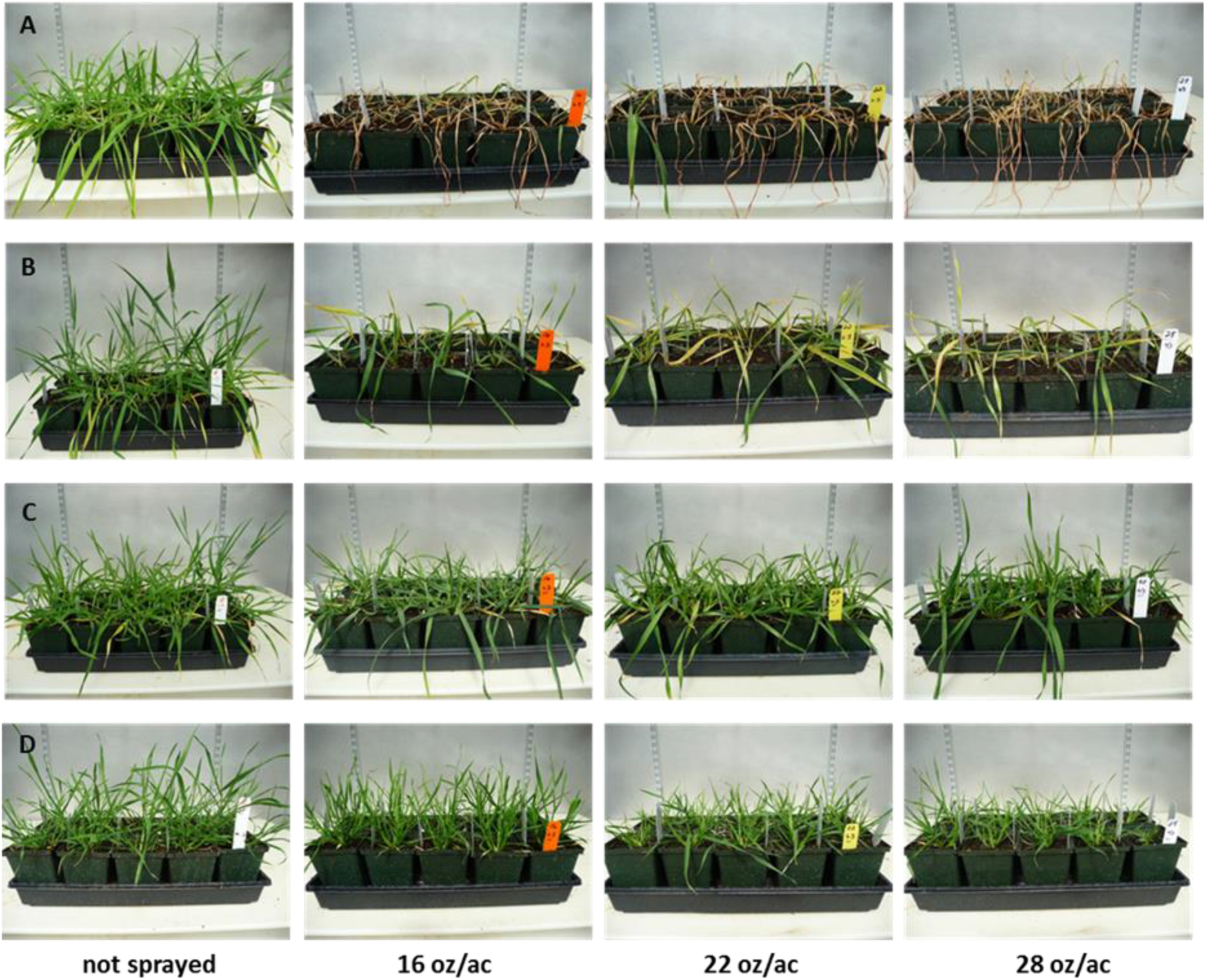
Greenhouse glyphosate spray chamber tests of different wheat genotypes. **A** represents the unmutagenized parental cultivar Express. **B** is the homozygous triple mutant line (genotype **B** in **Figure 3B**). C shows the genotype with the homozygous TIPS alleles on chromosomes 7A and 7D EPSPS homoeologs (genotype **C** in **Figure 3B**). **D** represents the genotype with TIPS mutations as well as the chromosome 4A T_102_I allele (genotype **D** in **Figure 3B**). Flats contained ~30 plants for each genotype except panel **B** whose flats contained 13 plants. Flats were sprayed at the 5-leaf stage with the indicated rate/acre equivalent of Roundup PowerMAX^®^. Images were taken two weeks after herbicide application.

**Figure 6.**
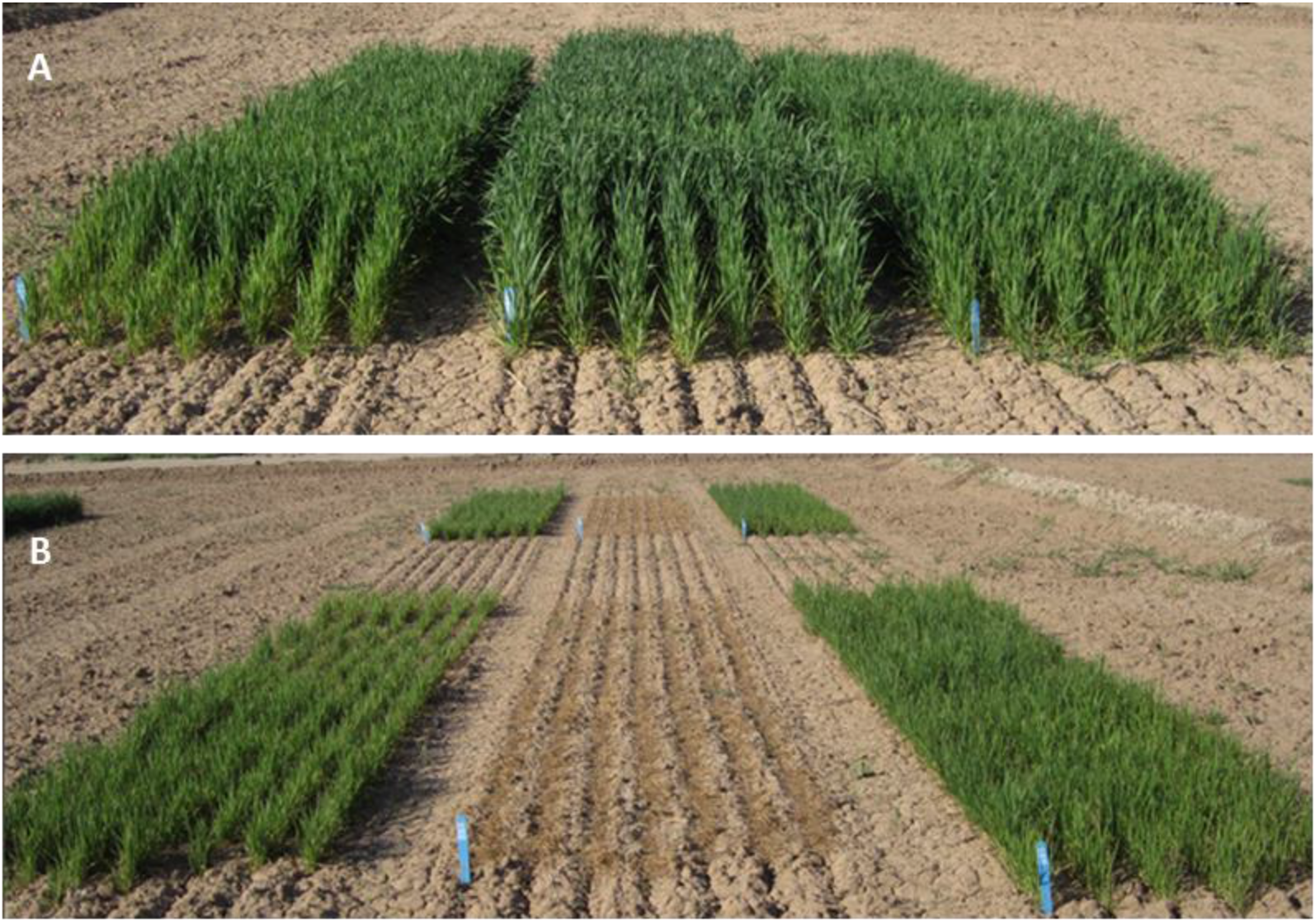
Field plots of wild type and TIPS allele containing lines without (A) and with (B) glyphosate herbicide treatment. In both panels **A** and **B**, the middle plot consists of the parental “Express” cultivar, while left and right plots are two separate lines containing chromosome 7A and 7D TIPS alleles (genotype **C** in **Figure 3B**). Panel **A** represents unsprayed plots, while in the foreground of panel **B**, plots were sprayed with the equivalent of 22 fluid ounces/acre of Roundup PowerMAX^®^. Plots in the background of panel **B** were sprayed with the equivalent of 16 fluid ounces/acre of Roundup PowerMAX^®^.

## Discussion

Here, we describe the creation, identification and development via TILLING of wheat lines that have substantially increased tolerance to the herbicidal effects of glyphosate compared to the parental cultivar, Express. These lines contain both chromosome 7A and 7D TIPS EPSPS alleles, while having a wild type EPSPS allele on chromosome 4A. The TIPS alleles, 11 nucleotides apart in the active site of this enzyme, render EPSPS insensitive to the effects of glyphosate and, similarly, plants containing these alleles have substantially increased resistance to this herbicide. Although others have generated wheat with partial glyphosate resistance, these wheat lines were identified through forward screens and the genetic basis of the resistance is not completely determined (Aramrak et al. 2018).

A feature of TILLING is the ability to estimate the probability of identifying a desired mutation based on the induced SNP frequency in the population and the population size. At the commencement of this project, with a TILLING population size of approximately 10,000 and a mutation frequency of 1/24kb (Slade et al. 2005), we estimated an 80% probability of identifying at least one of the desired mutations. In the event, we exceeded expectations, and one of the priority alleles, T_102_I on the 7A chromosome, was identified twice in independent wheat lines (**Figure 3A**).

An additional feature of mutagenic treatment with the DNA-alkylating chemical ethyl methanesulfonate (EMS), which was used in the creation of our wheat TILLING resource, is that approximately 95% of induced SNPs are G-C to A-T transitions (Greene et al. 2003). Thus, one can predict the spectrum of expected missense alterations in codons. In our work, we identified only two induced SNPs from more than one hundred sequenced mutations that were not expected EMS-induced G-C to A-T transitions, namely L_95_H in the 4A EPSPS homoeolog, and L_107_V in the 7D EPSPS homoeolog (**Figure 2**). While the two priority alleles, T_102_I and P_106_S, are expected missense changes, an additional allele that confers glyphosate insensitivity in some EPSPS enzymes, namely the missense G_101_A alteration (Dong et al. 2019) (Funke et al. 2006), is not expected from mutagenesis with EMS, and represents a limitation of this mutagen. The two mutations we did identify in this glycine (G_101_R on the 4A EPSPS homoeolog and G_101_E on the 7D homoeolog; **Figure 2**) are expected G-C to A-T EMS-induced transitions. The effects of these mutations on the EPSPS enzyme and on the plants containing them remain to be fully characterized.

This work is another illustration of the practical utility of TILLING and the remarkable ability of wheat to tolerate a high mutation load, which was also demonstrated by the hundreds of new alleles identified in the targeted genes in our earlier work (Moehs et al. 2019) (Slade et al. 2005) (Slade et al. 2012). TILLING in wheat has been applied by others for additional practical breeding efforts, such as generating lines resistant to the powdery mildew pathogen (Acevedo-Garcia et al. 2017). Aside from the particular result of glyphosate tolerance, our work also serves as an example of the ability to use TILLING to identify multiple induced SNPs nearby in the same gene after two rounds of mutagenesis (**Figure 2**). In addition to the TIPS alleles, we found other double mutants in EPSPS including one combination of mutants on the chromosome 7D homoeolog two amino acids apart (A_100_V, T_102_I, 6 nucleotides apart) and another double mutant combination, also on the 7D homoeolog, in adjacent amino acids (T_102_I, A_103_T) only 2 nucleotides apart!

While we have not conducted *in vitro* enzyme assays of the wheat EPSPS containing the TIPS mutations, the phenotypic effects on wheat plants containing these alleles in response to glyphosate treatment imply that in wheat, as in other tested plants (Dill 2005), these mutations render the enzyme insensitive to glyphosate. Nevertheless, evaluation of EPSPS enzymes with the TIPS allele in other species (Dong et al. 2019) (Yu et al. 2015) demonstrates that these TIPS enzymes, while having several thousand-fold elevated glyphosate inhibitory concentrations (K_i_), and near normal K_m_s for their natural substrate, phosphoenolpyruvate, also exhibited reductions in their V_max_, to a level of about 12% of the normal enzyme. The wheat TIPS EPSPS enzymes also have a lower V_max_ than the native enzyme. Suggestive evidence that this is the case comes from the fact that all plants having homozygous TIPS variants on the 7A and 7D homoeologs as well as the homozygous 4A T_102_I allele exhibited reduced vigor. This suggests both that the three homoeologous EPSPS genes are the only active gene copies encoding this enzymatic activity in wheat, and that wheat plants containing only mutant copies of this enzyme are impaired in their ability to supply the required products of the shikimate pathway. Additional experiments will be required to confirm this supposition. This result implies that one unmutated copy of EPSPS is required to minimize any potential fitness cost of these alleles and to maintain plant vigor in the absence of glyphosate treatment. Other possible combinations of novel EPSPS alleles including, for example, a wheat line with one TIPS and one wild type allele on either the 7A or 7D chromosome in combination with the chromosome 4A P_106_L mutation (**Figure 3A**) have yet to be characterized in detail.

The TILLING wheat lines we generated are not as resistant to glyphosate as transgenic glyphosate tolerant wheat (Zhou et al. 2003) since the transgenic wheat was developed using a strong constitutive promoter to express the glyphosate-insensitive EPSPS gene. Nevertheless, our non-transgenic glyphosate-tolerant wheat is resistant to commercial levels of glyphosate herbicide. Non-transgenic glyphosate tolerant wheat may have uses in weed and disease control (Anderson and Kolmer 2005) (Feng et al. 2005) and this wheat may benefit from advancements in robotic weeding technology that enables targeted herbicide spraying (Liu and Bruch 2020). In addition, the glyphosate tolerance may, in the future, be further improved by adding additional alleles in genes that generate glyphosate tolerance by independent mechanisms-e.g. reduced glyphosate translocation.

Although the advent of CRISPR/Cas9 and related gene editing methods to modify plant genomes has taken plant research by storm (Yin et al. 2017), and may supplant TILLING for some applications, the recent decision by the European Union to subject CRISPR-derived crops to regulatory requirements similar to transgenic crops (Callaway 2018) means that crops modified by TILLING have an easier path to market than CRISPR-derived crops in certain geographies. Finally, by expanding the conception of what can be achieved by conventional breeding, examples such as ours of identifying multiple missense mutations in the same gene may influence the discussion of the regulation of alleles derived by editing methods.

## Materials and Methods Plant Material

Hexaploid wheat (cv. Express) with mutations in the EPSPS gene in each of the 7A, 7D and 4A homoeologs were crossed to each other and to the parental cultivar. Different genetic classes were identified by KASP genotyping (https://www.biosearchtech.com/support/education/kasp-genotyping-reagents) using probes developed to specific SNPs and genomic DNA isolated from seedling leaf tissue. Wheat plants were sown in Sunshine Mix #3 and grown in pots in Conviron chambers and in the greenhouse as described by Slade et al. (Slade et al. 2012).

### TILLING and PCR

Approximately 10,000 individual M2 mutant genomic DNAs from spring wheat cultivar Express were screened. TILLING of wheat was conducted according to Slade et al., (2005) as follows: The M2 wheat DNA was pooled into groups of two individual plants. The DNA concentration for each individual within the pool was approximately 2 ng/μl with a final concentration of 4 ng/μl for the entire pool. Then, 5 μl of the pooled DNA samples (or 20 ng wheat DNA) was arrayed on microtiter plates and subjected to gene-specific PCR. Amplification and TILLING was performed exactly as described (Moehs et al. 2019).

### Assessment of glyphosate resistance

For growth chamber tests (**Figure 4**), wheat seeds were germinated in glass tubes with Phytagel medium containing or lacking 0.15mM glyphosate (Sigma-Aldrich). Plantlets were grown for two weeks in Conviron growth chambers under 18 hour illumination. For greenhouse spray chamber experiments (**Figure 5**), three week old seedlings mostly at the 5-leaf stage, grown in Sunshine Mix #3, were sprayed with a Teejet 9502EVS Nozzle at a 95 degree spray angle at 0.2 GPM (based on 40 psi), 21 inch above the edge of the flat. Roundup PowerMAX^®^ with 47.8% active ingredient was diluted to the appropriate equivalent fluid ounces/acre concentration. Images were taken at one week intervals post spray.

## Acknowledgements

For excellent technical assistance, the authors are grateful to Will Hines, Maricela Soltero de Torres and Steve Strickland. For his support of this project the authors thank Vic Knauf.

## Conflict of Interest Statement

Authors C.P.M, M.N.S., J.C.M. and A.J.S. are co-inventors on patents (US and Australia Patent Number 2016288257) related to the described research entitled: “Wheat having resistance to glyphosate due to alterations in 5-enol-pyruvylshikimate-3 phosphate synthase” and all authors were employed by Arcadia Biosciences, which has a commercial interest in the described research.

